# Suppression of autophagy induces senescence in the heart

**DOI:** 10.1101/2024.05.26.595978

**Authors:** Peiyong Zhai, Eun-Ah Sung, Yuka Shiheido-Watanabe, Koichiro Takayama, Yimin Tian, Junichi Sadoshima

**Affiliations:** Department of Cell Biology and Molecular Medicine, Cardiovascular Research Institute, Rutgers-New Jersey Medical School, 185 South Orange Ave, Newark, NJ 07103

**Keywords:** Atg7, Doxorubicin, Cardiac dysfunction, Senolysis, ABT-263, Beclin 1

## Abstract

Aging is a critical risk factor for heart disease, including ischemic heart disease and heart failure. Cellular senescence, characterized by DNA damage, resistance to apoptosis and the senescence-associated secretory phenotype (SASP), occurs in many cell types, including cardiomyocytes. Senescence precipitates the aging process in surrounding cells and the organ through paracrine mechanisms. Generalized autophagy, which degrades cytosolic materials in a non-selective manner, is decreased during aging in the heart. This decrease causes deterioration of cellular quality control mechanisms, facilitates aging and negatively affects lifespan in animals, including mice. Although suppression of generalized autophagy could promote senescence, it remains unclear whether the suppression of autophagy directly stimulates senescence in cardiomyocytes, which, in turn, promotes myocardial dysfunction in the heart. We addressed this question using mouse models with a loss of autophagy function.

Suppression of general autophagy in cardiac-specific *Atg7* knockout (*Atg7*cKO) mice caused accumulation of senescent cardiomyocytes. Induction of senescence via downregulation of *Atg7* was also observed in chimeric *Atg7* cardiac-specific KO mice and cultured cardiomyocytes *in vitro*, suggesting that the effect of autophagy suppression upon induction of senescence is cell autonomous. ABT-263, a senolytic agent, reduced the number of senescent myocytes and improved cardiac function in *Atg7*cKO mice. Suppression of autophagy and induction of senescence were also observed in doxorubicin-treated hearts, where activation of autophagy alleviated senescence in cardiomyocytes and cardiac dysfunction. These results suggest that suppression of general autophagy directly induces senescence in cardiomyocytes, which in turn promotes cardiac dysfunction.

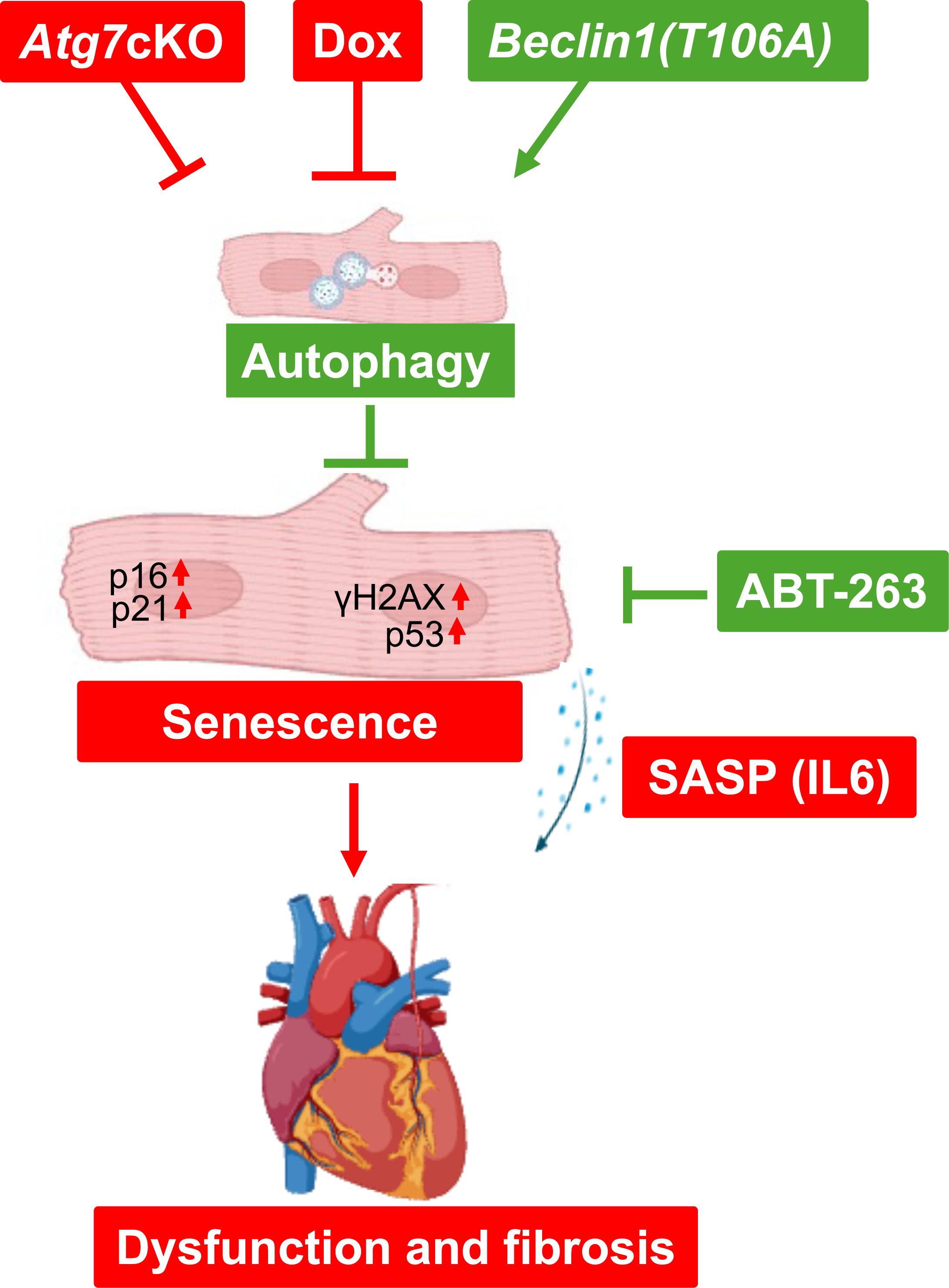

## 1. Introduction

Aging is a significant risk factor promoting heart failure [1, 2]. Mitochondrial dysfunction is commonly observed in aging hearts [3], and increases in oxidative stress, DNA damage and inflammation drive aging in the heart [4]. Oxidative stress promotes protein misfolding and aggregation, which exacerbates organelle dysfunction and cell death [5].

Autophagy is a major mechanism of cellular degradation that it degrades protein aggregates and dysfunctional organelles [6]. Autophagy is generally *inhibited during aging* [5, 7, 8], whereas stimulation of autophagy extends lifespan in mice [9]. Autophagy protects against aging by mediating protein and organelle quality control mechanisms [5] [10]. On the other hand, suppression of autophagy impairs cellular homeostasis and leads to organ dysfunction and aging. However, the underlying mechanisms remain to be elucidated [5, 8].

Cellular senescence is a condition in which cells limit their own replicative potential while inducing reprogramming or aging of neighboring cells through paracrine mechanisms [11]. Thus, senescence is a major driver of organ aging. Since senescence is induced primarily through telomere attrition in proliferating cells, whether senescence occurs in adult cardiomyocytes has been controversial. However, telomere attrition can take place together with DNA damage in failing hearts [12, 13]. Length-independent telomere damage caused by aging [14] also induces senescence in terminally differentiated cardiomyocytes (CMs) [15–19]. In general, autophagy negatively regulates senescence by maintaining cellular homeostasis [10, 20] or by degrading positive regulators of senescence, such as GATA4 [21]. However, autophagy also promotes senescence under some conditions. Amino acids and other metabolites recycled through autophagy support the production of SASPs [22, 23]. Autophagy also promotes senescence by degrading specific substrates [24] or by promoting survival of senescent cells in chronic kidney disease [25]. In addition, coordinated activation of selective autophagy is necessary for maintenance of the senescent state in fibroblasts [26]. Thus, *how autophagy affects senescence appears to be context-dependent* [24, 25, 27–34].

We reasoned that *how autophagy affects cardiomyocyte senescence* may provide clues to elucidating how aging is promoted in the heart. We here tested our hypothesis that *downregulation of general autophagy facilitates cardiac aging through induction of autophagy in cardiomyocytes*. We investigated 1) whether suppression of autophagy in Atg7 cardiac-specific knockout (*Atg7*cKO) mice and the mouse model of doxorubicin (Dox)-induced cardiomyopathy induces senescence in cardiomyocytes and 2) the causative involvement of cellular senescence in the development of cardiomyopathy.

## 2. Materials and Methods

### 2.1 Genetically modified mice

Cardiac-specific homozygous *Atg7* knockout mice (*Atg7*cKO) were generated by crossing *Atg7* flox homozygous (fl/fl) mice with αMHC-Cre mice [35, 36]. *Beclin 1(T106A)* knockin mice were generated by the Genome Editing Shared Resource at Rutgers, The State University of New Jersey. All animal experiments were approved by the Institutional Animal Care and Use Committee of the New Jersey Medical School, Rutgers, The State University of New Jersey. The animal use in this study also conformed to the Guide for the Care and Use of Laboratory Animals published by the US National Institutes of Health (8th edition, 2011). All animal experiments used both male and female mice. The investigators were blinded to the genotype groups during the experiments.

### 2.2 Doxorubicin (Dox) administration to mice

Doxorubicin (Sigma) was administered to mice via intraperitoneal injection at a dose of 5 mg/kg (dissolved in saline) once a week for 4 consecutive weeks [37]. Mice injected with saline were used as controls.

### 2.3 ABT-263 (Navitoclax) administration to mice

ABT-263 (Navitoclax, Selleckchem), dissolved in ethanol/polyethylene glycol 400/Phosal 50 PG at a ratio of 10:30:60, was administered to mice through oral gavage feeding at 50 mg per kg body weight per day (mg/kg/d) for 5 days per cycle for two cycles with a 1-week interval between the cycles [38]. Mice fed the solvent were used as controls.

### 2.4 AAV9-cTnt-Cre-GFP injection into mice

The mice were anesthetized with 1.5% isoflurane. Under surgical plane of anesthesia, AAV9*-cTnt-Cre-GFP* or a control AAV9, at the dose of 1-2x10^11^ vector genomes/mouse, was injected into the mice via the external jugular vein as described previously [39].

### 2.5 Primary culture of neonatal rat ventricular cardiomyocytes

Primary cultures of ventricular cardiomyocytes were prepared from 1-day-old Charles River Laboratories (Crl)/Wistar Institute (WI) BR-Wistar rats (Harlan Laboratories, Somerville, NJ, USA) and maintained in culture as described previously [40]. A cardiomyocyte-rich fraction was obtained by centrifugation through a discontinuous Percoll gradient.

### 2.6 Senescence associated-β-galactosidase (SA-β-gal) staining of cardiomyocytes and cardiac tissue sections

Neonatal rat ventricular myocytes (NRVMs) that were treated with PBS or Dox or transduced with adenovirus harboring shRNA *scramble* (sh*Ctrl*) or shRNA *Atg7* (sh*Atg7*) were stained with the SA-β-gal staining kit (Cell Signaling Technologies, #9860), according to the protocol provided by the manufacturer. To evaluate the SA-β-gal in cardiac tissue, frozen cardiac tissue sections were prepared and stained with the SA-β-gal staining kit according to the protocol provided by the manufacturer.

### 2.7 Echocardiography

Transthoracic echocardiography was conducted using a high-resolution Micro-Ultrasound system (Vevo 3100, FUJIFILM VisualSonics Inc., Toronto, Canada). Two-dimension (short-axis)-guided M-mode measurements (at the level of the papillary muscles) of the left ventricle (LV) internal diameter were taken from 3 or more beats and averaged. LV end-diastolic dimension (LVEDD) was measured at the time of the apparent maximal LV diastolic dimension, whereas LV end-systolic dimension (LVESD) was measured at the time of the most anterior systolic excursion of the posterior wall. LV ejection fraction (LVEF) and LV fractional shortening (LVFS) were analyzed using Vevo LAB software (FUJIFILM VisualSonics, Inc.) to evaluate systolic function.

### 2.8 Immunoblotting analysis

Cardiac tissue homogenates were prepared from the LV apex. Cell lysates were prepared from primary cultures of rat ventricular myocytes 24-72 hours after adenovirus infection. Both the homogenates and the cell lysates were prepared in a radioimmunoprecipitation assay (RIPA) buffer containing 150 mM NaCl, 1% Triton-X 100, 0.5% sodium deoxycholate, 0.1% sodium dodecyl sulphate, and 50 mM Tris, pH 8.0, and supplemented with a protease inhibitor cocktail (Sigma), 5 mM NaF, and 1 mM sodium orthovanadate. We used antibodies against phospho-H2AX (S139) (CST, #9718), IL-6 (Thermo Fisher Scientific, #P620), P16 (SCBT, #SC-1661), p19 (Abcam, #ab80), Atg7 (CST, #8558S), P21 (SCBT, #SC-6246), P53 (Bioss, #8687R), GAPDH (CST, #2118), LC3 (MBL, #M186-3), and a-tubulin (Sigma, #T6199).

### 2.9 Evaluation of autophagic flux

To evaluate autophagic flux *in vivo,* chloroquine (CQ) was injected (10 μg/kg) intraperitoneally [41]. Three to four hours later, mice were euthanized and heart homogenates were prepared. The amount of LC3 was evaluated with immunoblot analyses with an anti-LC3 antibody. To evaluate autophagic flux *in vitro*, NRVMs were transduced with GFP-LC3 adenovirus with or without Dox treatment for 48 hours. Bafilomycin A1 (Baf A1) was applied for three hours before cell fixation. The number of GFP-LC3 puncta per cell was counted.

### 2.10 Immunofluorescence analysis

Mouse hearts were harvested, fixed with 10% formalin and embedded in wax and cross-sections (5-μm thick) were prepared. Neonatal cardiomyocytes were cultured on coverslips and fixed with 4% PFA. An overnight incubation with specific antibodies (P16, IL-6 or phospho-H2AX (S139) and cTNT (Invitrogen, #MA-5-12960)) was followed by 2 hours of incubation with secondary antibody conjugated with Alexa Fluor 488, 555, 594 or 647 dye (Life Technologies). Samples were washed and mounted with a reagent containing DAPI (VECTASHIELD; Vector Laboratories). The analyses were performed in a blinded manner by fluorescence microscopy.

For analysis of γH2AX-positive cardiomyocytes, the total number of γH2AX-positive nuclei in cardiomyocytes in the whole section was counted. The total number of cardiomyocytes in the whole section was estimated as the product of the average number of cardiomyocytes per image and the total number of images per section. The percentage of γH2AX-positive cardiomyocytes was calculated by the equation: the number of γH2AX-positive cardiomyocytes/total number of cardiomyocytes in the section multiplied by 100.

### 2.11 Cytokine array analysis

NRVMs were transduced with sh*Ctrl* or sh*Atg7* for 48 hours and the supernatants of the cultures were collected for cytokine array analysis using a rat cytokine array kit (Abcam, # ab133992) according the protocol provided by the manufacturer.

### 2.12 Statistical analysis

The data are presented as mean±SEM and were first analyzed using analysis of variance (ANOVA). If significant differences were observed, post-hoc analysis was performed using the t test with Bonferroni correction. A p value less than 0.05 was considered significant.

## 3. Results

### 3.1 The *Atg7*cKO mouse heart exhibits more senescence

We have shown previously that autophagy is suppressed completely in cardiac-specific homozygous *Atg7* knockout (KO) (*Atg7*cKO) mouse hearts [35, 36]. We therefore used *Atg7*cKO mice as a model of autophagy downregulation in cardiomyocytes to study how inhibition of autophagy affects senescence in cardiomyocytes. Cardiac function in *Atg7*cKO mice was not significantly different from that in wild-type (WT) mice at 2-4 months of age [36]. Atg7 in the heart was downregulated in *Atg7*cKO mice compared to in WT mice. Markers of senescence, including cell cycle inhibitors (P16 and P21), P53, a DNA damage marker (γH2AX), and an inflammatory cytokine (IL-6), were upregulated in the *Atg7*cKO mouse heart compared to in the WT mouse heart (**Fig. 1**). Double staining of cardiac sections with P16, γH2AX or IL-6 and cardiac troponin T (cTNT), a cardiomyocyte marker, showed that there were more P16-, IL6- and γH2AX-positive cardiomyocytes in *Atg7*cKO than in WT mouse hearts (**Fig. 2**). Quantitative analyses showed that P16-positive cells were primarily cardiomyocytes rather than non-myocytes. Frozen sections of the heart also showed that *Atg7*cKO mouse hearts have a greater level of senescent activated β-galactosidase positivity than WT mouse hearts (**Supplemental Fig. 1**). These results suggest that senescence markers are upregulated in *Atg7*cKO mouse hearts at 2-4 months of age compared to in WT mouse hearts.

**Figure 1.**
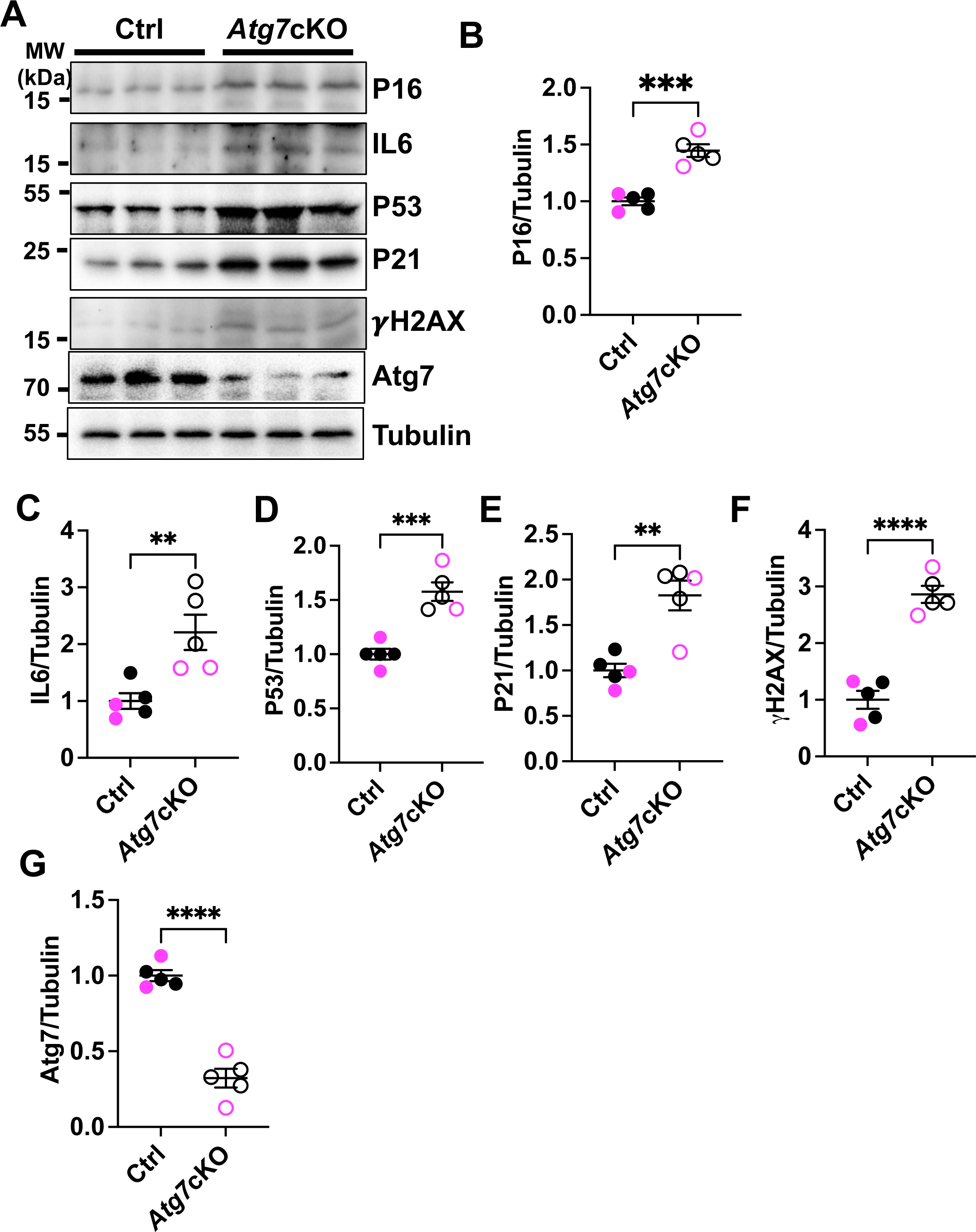
*Atg7*cKO mouse hearts have higher levels of cellular senescence markers. A. Immunoblotting of P16, IL6, P53, P21, γH2AX, Atg7, and tubulin in *Atg7*fl/fl (Ctrl) and *Atg7*cKO mouse hearts. B-G. Relative band density of the cellular senescence markers shown in A. n (the number of mice) =5 in each group. Black dots/circles denote data from male mice and pink dots/circles denote data from females. Data are presented as Mean±SEM. Statistical analysis was conducted using Prism unpaired t test. ** P<0.01, *** P<0.001, **** P<0.0001. P<0.05 was considered statistically significant.

**Figure 2.**
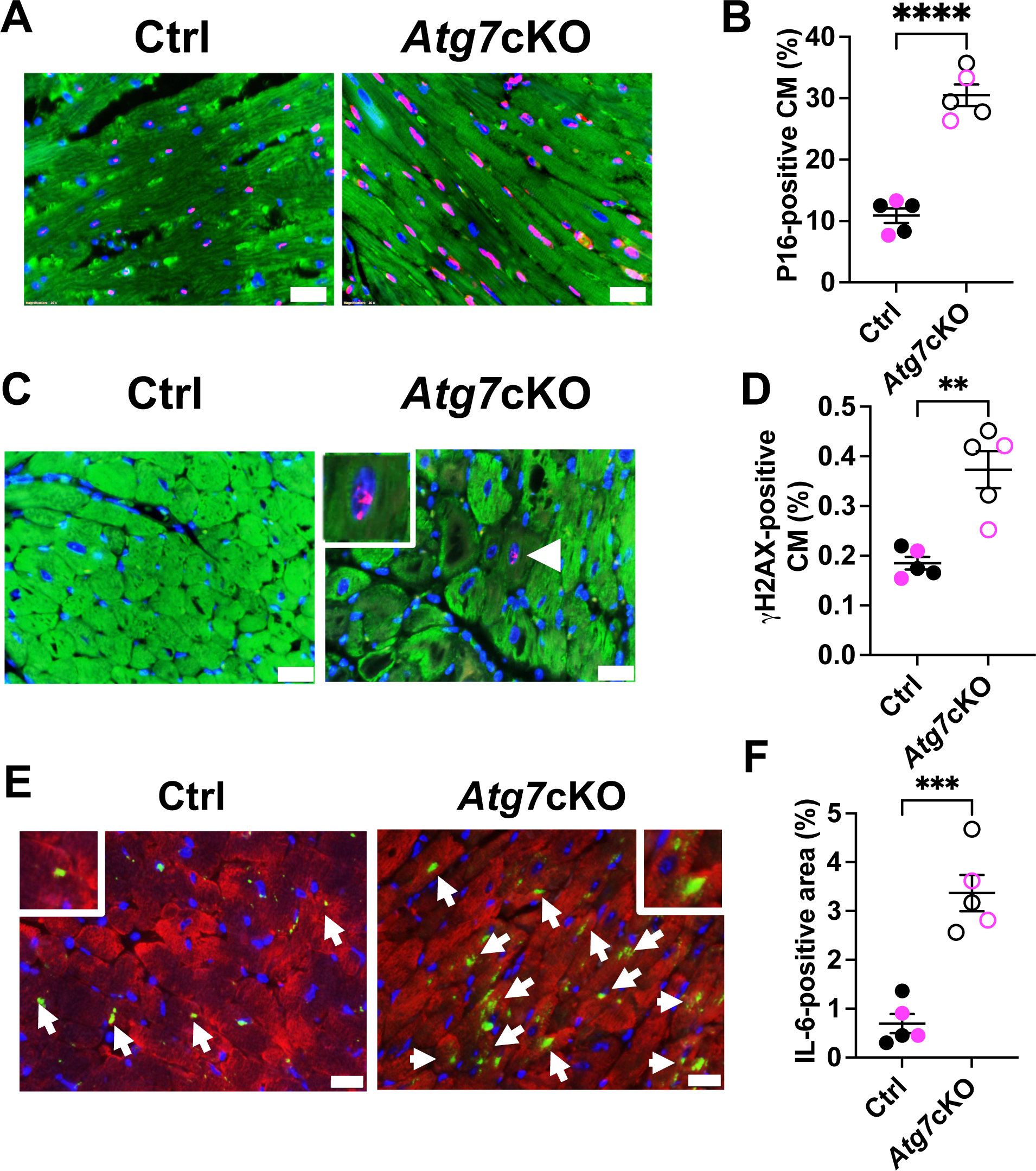
The *Atg7*cKO mouse heart exhibits more senescence. A. P16 (red) and cTNT (green) immunostaining. B. Percentage of P16-positive cardiomyocytes. C. γH2AX (red) and cTNT (green) immunostaining. D. Percentage of γH2AX-positive cardiomyocytes. E. IL6 (green) and cTNT (red) immunostaining. F. Percentage of IL6-positive area. In A, C and E, scale bars = 20 μm. Arrow heads point to positive nuclei. In B, D and F, n (the number of animals) =5. Data are presented as Mean±SEM. Black dots/circles show data from male mice and pink dots/circles show data from females. Statistical analysis was conducted using Prism unpaired t test. **** P<0.0001, *** P<0.001, ** P<0.01. P<0.05 was considered statistically significant.

### 3.2 Chimeric downregulation of Atg7 in the mouse heart induces senescence in cardiomyocytes

In order to assess the more direct effect of autophagy suppression upon cardiomyocyte senescence, we generated genetically altered mice in which Atg7 is downregulated only in a few cardiomyocytes via jugular vein injection of AAV9-*cTnt-Cre-Gfp* into *Atg7flox/flox* mice. The goal in this experiment was to compare the level of senescence in cardiomyocytes with Atg7 downregulation and those without Atg7 downregulation in the same heart. As expected, we observed only a small reduction of left ventricular (LV) function in the presence of AAV9-*cTnt-Cre-Gfp* (**Fig. 3A-C** and Supplemental Table 1). In this condition, P16-positivity was observed primarily in Cre-GFP positive cells, rather than in Cre-GFP negative cells, in the *Atg7flox/flox* mice injected with AAV9-*cTnt-Cre-Gfp* (**Fig. 3D-F**). This result suggests that Atg7 downregulation induces senescence in cardiomyocytes in a cell autonomous manner in the heart *in vivo*.

**Figure 3.**
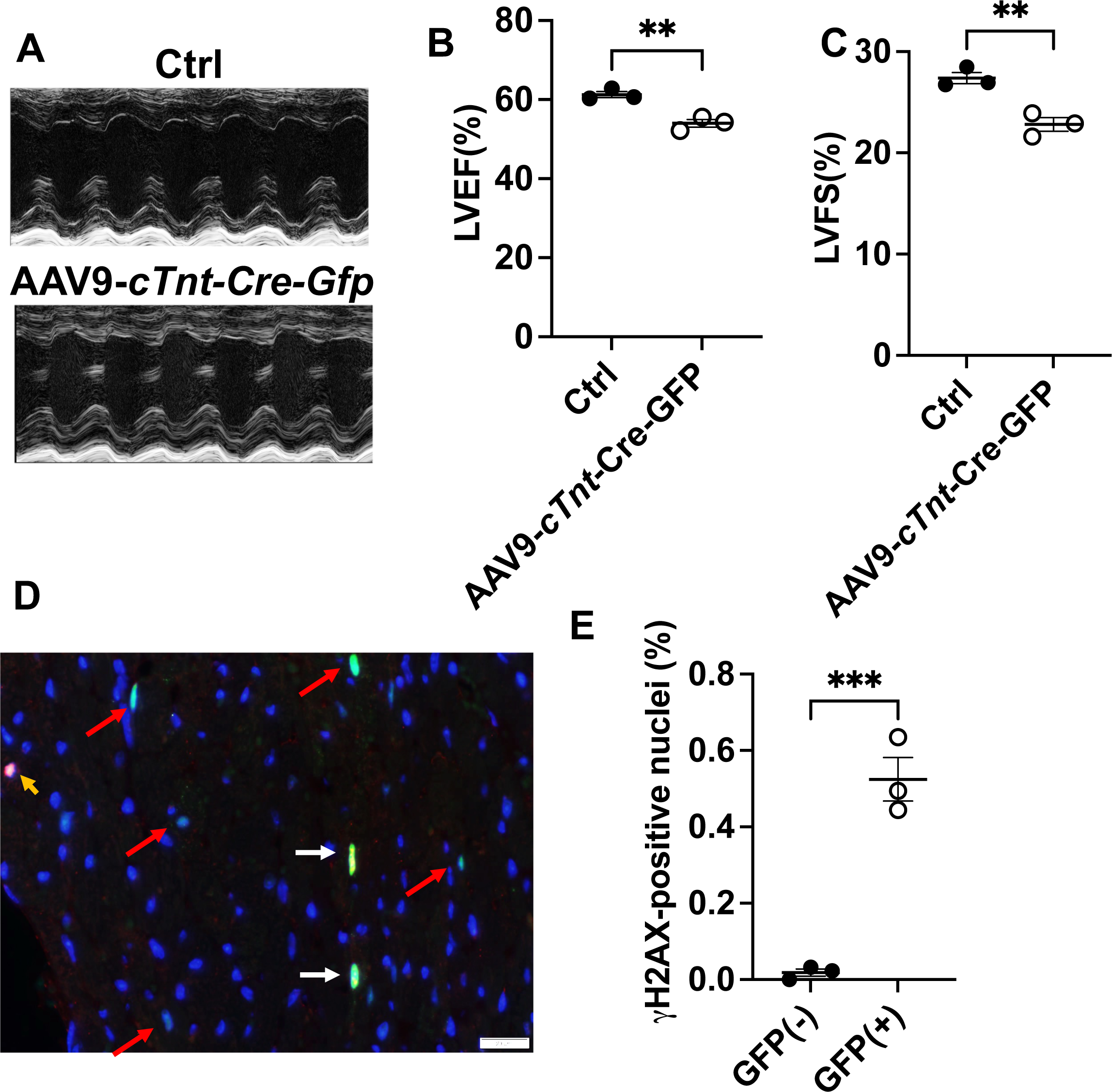
Chimeric downregulation of *Atg7* in cardiomyocytes in mice induced senescence. *Atg7*fl/fl mice were injected with AAV9-cTnt-Cre-GFP (GFP) or a control AAV9 (Ctrl). Hearts were harvested 4 weeks after AAV injection and processed for immunostaining. A. Representative M-mode echocardiographic images. B. Left ventricular ejection fraction (LVEF) in AAV9-cTnt-Cre-GFP-transduced or control AAV9-transduced mice. C. Left ventricular fractional shortening (LVFS) in AAV9-cTnt-Cre-GFP-transduced or control AAV9-transduced mice. In B-C, ** P<0.01. D. An image of γH2AX (red) immunofluorescence staining and GFP (green). White arrows point to γH2AX- and GFP-positive nuclei. Red arrows point to GFP-positive nuclei. Orange arrow points to γH2AX-positive nucleus. Scale bars = 20 μm. E. Percentage of γH2AX-positive nuclei in GFP-positive (GFP(+)) and GFP-negative (GFP(-)) cells. *** P<0.001. In B, C and E, n (the number of mice) =3. Data are presented as Mean±SEM. Statistical analyses were conducted using Prism unpaired t test. P<0.05 was considered statistically significant.

### 3.3 Downregulation of autophagy promotes senescence in cultured cardiomyocytes

To examine whether induction of senescence in cardiomyocytes by autophagy inhibition is cell-autonomous, we downregulated Atg7 in cultured neonatal rat ventricular cardiomyocytes (NRVMs) and evaluated the induction of senescence markers. As expected, treatment of NRVMs with shRNA *Atg7* downregulated Atg7. Downregulation of Atg7 was sufficient to upregulate senescence markers, including P16, IL-6, P53, P21, γH2AX, and SA-β-gal staining (**Fig. 4**), suggesting that induction of senescence in cardiomyocytes via suppression of autophagy is cell-autonomous. siRNA against Atg7 also induced senescence markers in NRVMs, confirming the validity of *Atg7* knock-down experiments (**Supplemental Fig. 2**). Cell senescence is also characterized by secretion of cytokines, termed the senescence-associated secretory phenotype (SASP). Thus, we conducted multiplex ELISA assays and found that Atg7 downregulation upregulated (>1.4 fold) CINC-2a [19], IL-6 [19], B7-2/CD86, MIP-3a [42], RAGE [43] and TNF-α [19] and downregulated (<0.8 fold) CNTF [44],[45], IL-1R6 [46], IL-2 [47], and CXCL5 [48] in NRVMs (Supplementary Table 2). These results suggest that suppression of autophagy upregulates proinflammatory cytokines while downregulating some cytokines known to be protective, consistent with the notion that suppression of autophagy in cardiomyocytes promotes the SASP phenotype.

**Figure 4.**
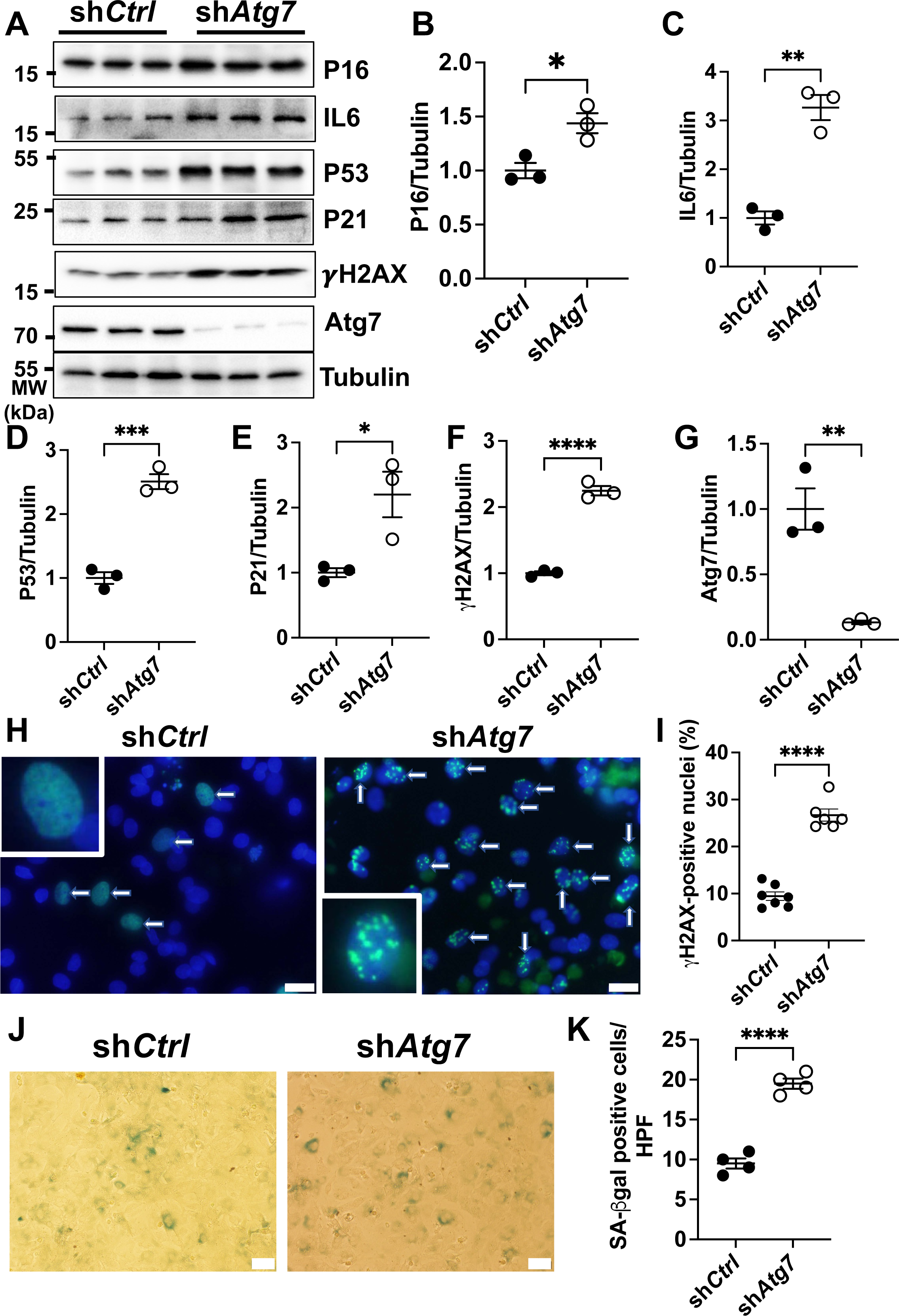
Reduction in *Atg7* promotes senescence in cultured cardiomyocytes. A. Cellular senescence markers (P16, IL6, P53, P21, and γH2AX), Atg7, and tubulin in neonatal rat ventricular myocytes (NRVMs) transduced with short-hairpin RNA against *Atg7* (sh*Atg7*) or a control short-hairpin RNA (shCtrl). B-G. Relative band density of the molecules shown in A. * P<0.05, ** P<0.01, *** P<0.001, **** P<0.0001. H. γH2AX immunofluorescence staining in NRVMs infected with sh*Atg7* or shCtrl. Whie arrows point to γH2AX-positive nuclei. Scale bars = 20 μm. I. Percentage of γH2AX-positive nuclei in NRVMs transduced with sh*Atg7* or shCtrl. *** P<0.005. J. Senescence associated (SA) β-galactosidase (β-gal) staining in NRVMs transduced with shCtrl or sh*Atg7*. Scale bars = 50 μm. K. Number of positive β-gal cardiomyocytes per high power field (HPF). **** P<0.0001. In this figure, n (the number of experiments) =3. Data are presented as Mean±SEM. The statistical analyses were conducted using Prism unpaired t test. P<0.05 was considered statistically significant.

### 3.4 Senescence promotes cardiac dysfunction in *Atg7*cKO mice

In order to investigate the functional significance of the induction of senescence in the *Atg7*cKO mouse heart, we treated *Atg7*cKO and WT mice at 6 months of age with either ABT-263, a senolytic [38, 49], or vehicle, as indicated in **Fig. 5A**. Senescence in cardiomyocytes, evaluated with γH2AX and cTNT staining, was decreased in the presence of ABT-263 in both *Atg7*cKO and WT mice, suggesting that ABT-263 successfully eliminated senescent cardiomyocytes (**Fig. 5BC**). ABT-263 treatment significantly reduced the number of P16-positive cardiomyocytes whereas it increased the number of P16- and TUNEL-double positive cardiomyocytes in *Atg7*cKO mouse hearts, suggesting that ABT-263 induces apoptosis in senescent cardiomyocytes particularly in *Atg7*cKO mouse hearts (**Supplemental Figure 3**). ABT-263 treatment also attenuated the increase in P16- and P21-positive cardiomyocytes in *Atg7*cKO mice (**Fig. 5D-G**). These results suggest that ABT-263 treatment effectively reduced cellular senescence in *Atg7*cKO mouse hearts. LV ejection fraction (LVEF) was significantly decreased in *Atg7*cKO mice at 6 months of age compared to in WT mince under vehicle-treated conditions, consistent with our previous results, suggesting that long-term suppression of autophagy induced LV dysfunction. ABT-263 treatment significantly improved LVEF in *Atg7*cKO mouse hearts (**Fig. 5HI** and Supplementary Table 3), suggesting that senolytic treatment improves LV function in *Atg7*cKO mice. Furthermore, our results suggest induction of cardiomyocyte senescence plays a causative role in mediating LV dysfunction in *Atg7*cKO mice.

**Figure 5.**
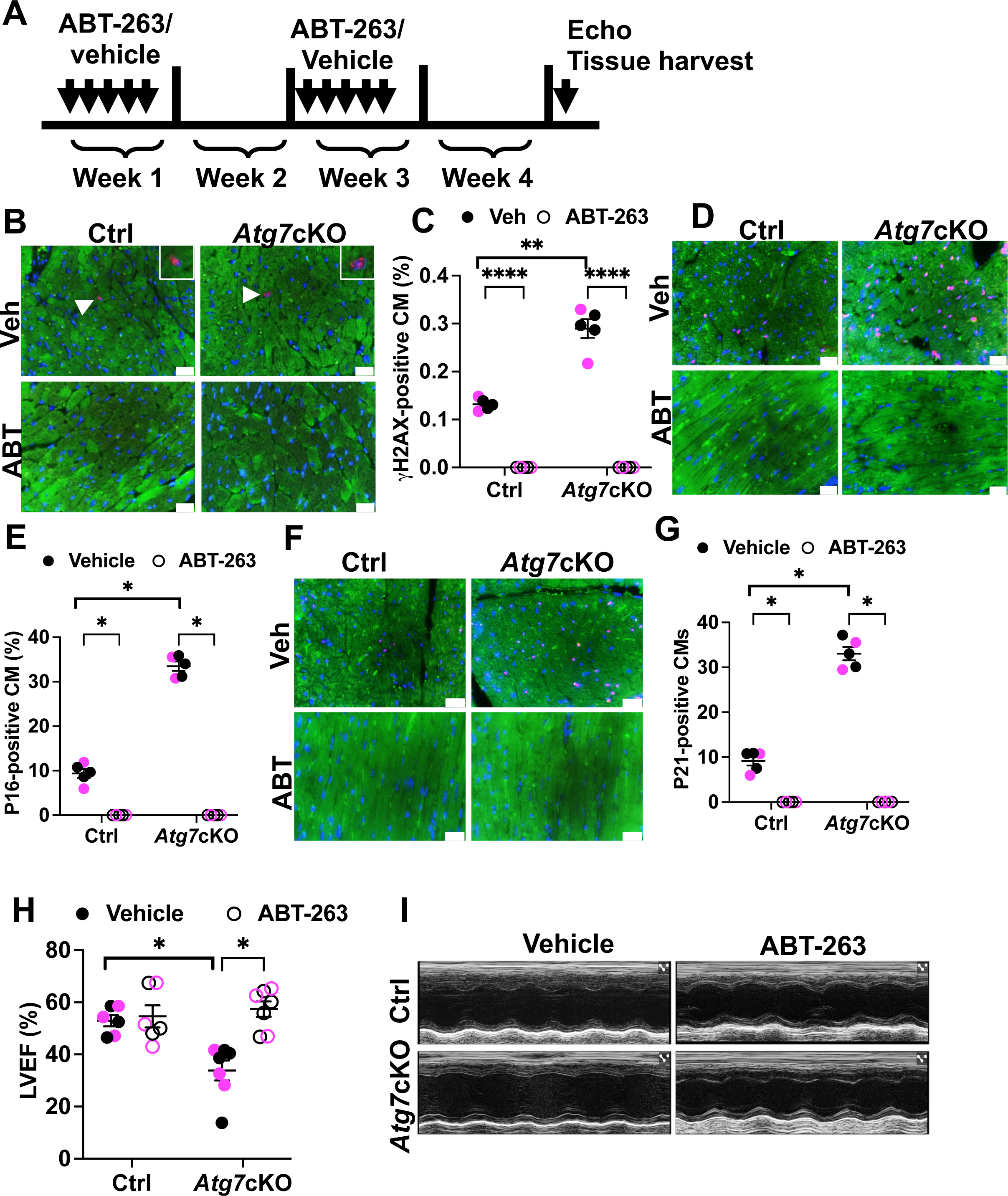
Senolysis improves cardiac function in *Atg7*cKO mice. A. A scheme showing the protocol of senolysis with ABT-263 in control (Ctrl) and *Atg7*cKO mice. In this experiment, we used mice with 6 months of age. B. Images of γH2AX (red) and cTNT (green) immunostaining in Ctrl and *Atg7*cKO mice. C. Percentage of γH2AX-positive cardiomyocytes in Ctrl and *Atg7*cKO mice. ** P<0.01. **** P<0.0001. D. Images of P16 (red) and cTNT (green) immunostaining in Ctrl and *Atg7*cKO mice. E. Percentage of P16-positive cardiomyocytes in Ctrl and *Atg7*cKO mice. * P<0.05. F. Images of P21 (red) and cTNT (green) immunostaining in Ctrl and *Atg7*cKO mice. G. Percentage of P21-positive cardiomyocytes. * P<0.05. H. Left ventricular ejection fraction (LVEF) in Ctrl and *Atg7*cKO mice treated with ABT-263 or vehicle. * P<0.05. I. Representative M-mode echocardiographic images. In this figure, n (the number of animals) =6 in each group. In C, E, G and H, black dots/circles show data from male mice and pink dots/circles show data from females. Data are presented as Mean±SEM. Statistical analyses were conducted using Prism 2-way ANOVA followed by multiple comparison t test and/or Bonferroni’s t test. P<0.05 was considered significant.

### 3.5 Doxorubicin (Dox)-induced cardiomyopathy is accompanied by the suppression of autophagy and induction of senescence

In order to further examine the significance of autophagy suppression in cardiomyocyte senescence, we used the mouse model of Dox-induced cardiomyopathy. We used a well-established protocol to induce cardiomyopathy in mice with Dox (5 mg/kg, cumulative dosage of 20 mg/kg, by once-per-week intraperitoneal injections) [37] (**Fig. 6A**). This dosage was selected based on the comparable pharmacokinetics of plasma Dox after a single dose of 5 mg/kg in mice to that observed in patients following a standard dose of Dox treatment (60 mg/m^2^).

**Figure 6.**
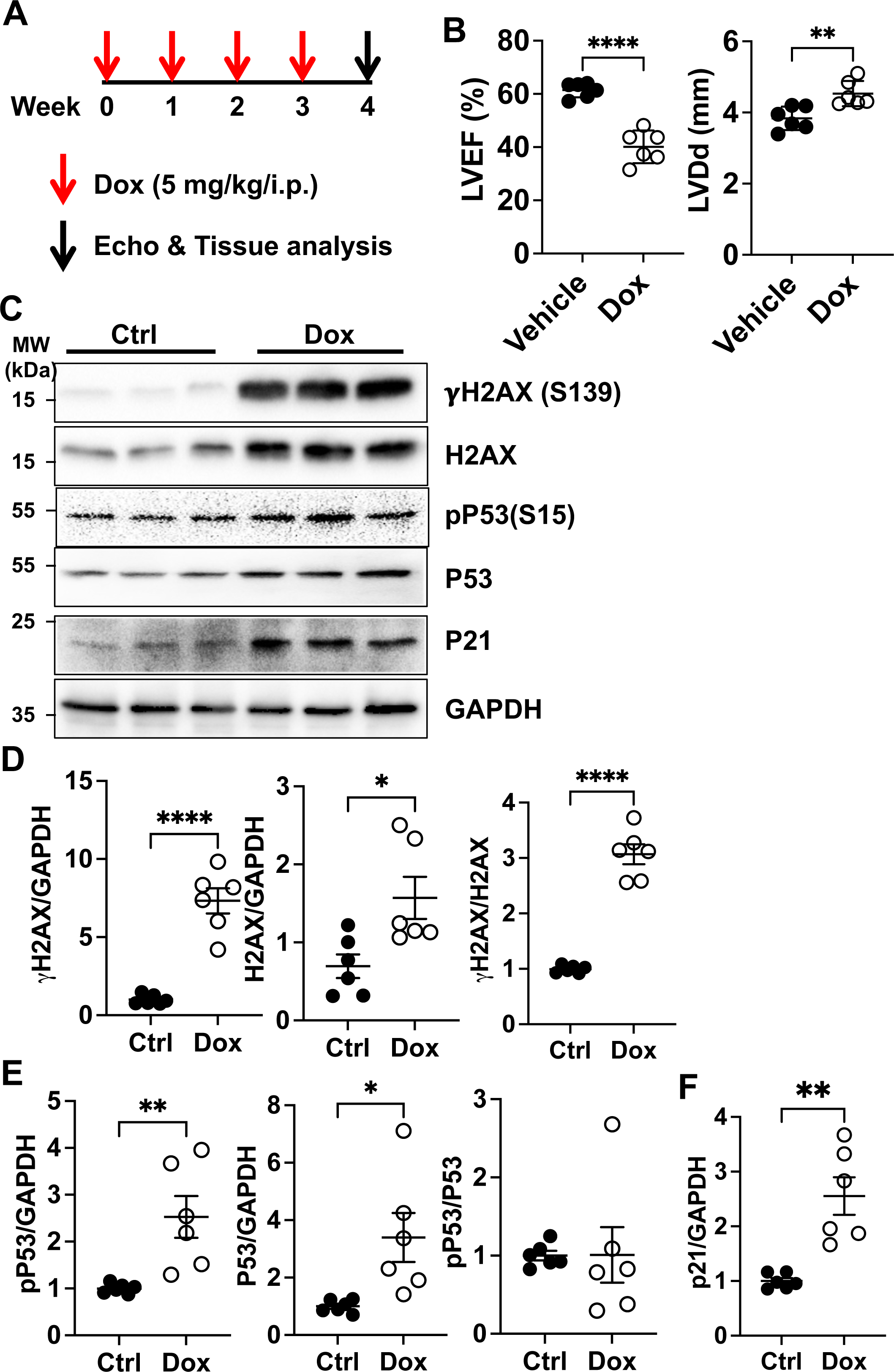
Doxorubicin (Dox) induced cardiomyocyte senescence both *in vitro* and *in vivo*. A. The Dox administration protocol. B. Echocardiographic (echo) analysis of ejection fraction (EF) and end-diastolic left ventricular internal dimensions (LVIDd). ** P<0.01, **** P<0.0001. C. Immunoblots of γH2AX, H2AX, pP53 (S15), P53, P21, and GAPDH in PBS-(Ctrl) or Dox-treated NRVMs. D. Relative band density of γH2AX and H2AX. E. Relative band density of pP53 and P53. F. Relative band density of P21. In D-F, * P<0.05, ** P<0.01, *** P<0.001, **** P<0.0001. Data are presented as Mean±SEM. The statistical analyses were conducted using Prism unpaired t test. P<0.05 was statistically significant.

Echocardiographic analyses showed that the Dox treatment induced decreases in LV contraction and increases in LV chamber size (**Fig. 6B**). Dox also induced upregulation of *p16*, *p21, IL-6, IL-1β*, and *IL-10* mRNA in LV homogenates (**Supplemental Fig. 4A**), suggesting that Dox induces cardiomyocyte senescence in the mouse heart. Dox (100 nM) also increased γH2AX (**Supplemental Fig. 4B**) and SA-β-gal positivity as evidenced by staining of cultured NRVMs (**Supplemental Fig. 4C**). Dox increased H2AX, γH2AX, and γH2AX/H2AX in cultured NRVMs as evaluated with immunoblot analyses (**Fig. 6CD**). Dox also increased P53, Serine 15 phosphorylated of P53 (pP53) (**Fig. 6CE**) and P21 (**Fig. 6CF**) in NRVMs. These results suggest that Dox induces senescence in cardiomyocytes in a cell-autonomous manner.

We investigated how Dox affects autophagy in the heart. To assess the autophagic flux, it is crucial to measure the degradation of LC3II in a lysosome-dependent manner. To evaluate the lysosome-dependent degradation, the lysosomal enzyme inhibitor, Chloroquine (CQ), was injected into mice 4 hours before euthanasia. Dox increased LC3II in the mouse heart compared to vehicle treatment (**Fig. 7A**). CQ injection increased the accumulation of LC3II in both Dox- and vehicle-treated hearts but the CQ-induced increase in LC3II was smaller in Dox-treated hearts than in control hearts (**Fig. 7A**), suggesting that autophagic flux is inhibited by the Dox treatment, consistent with previous studies [50]. Similar results were obtained in NRVMs treated with Dox (**Fig. 7B**), suggesting that Dox suppresses autophagic flux in cardiomyocytes in a cell-autonomous manner. Thus, like Atg7 downregulation, Dox treatment inhibits autophagy and induces senescence in cardiomyocytes.

**Figure 7.**
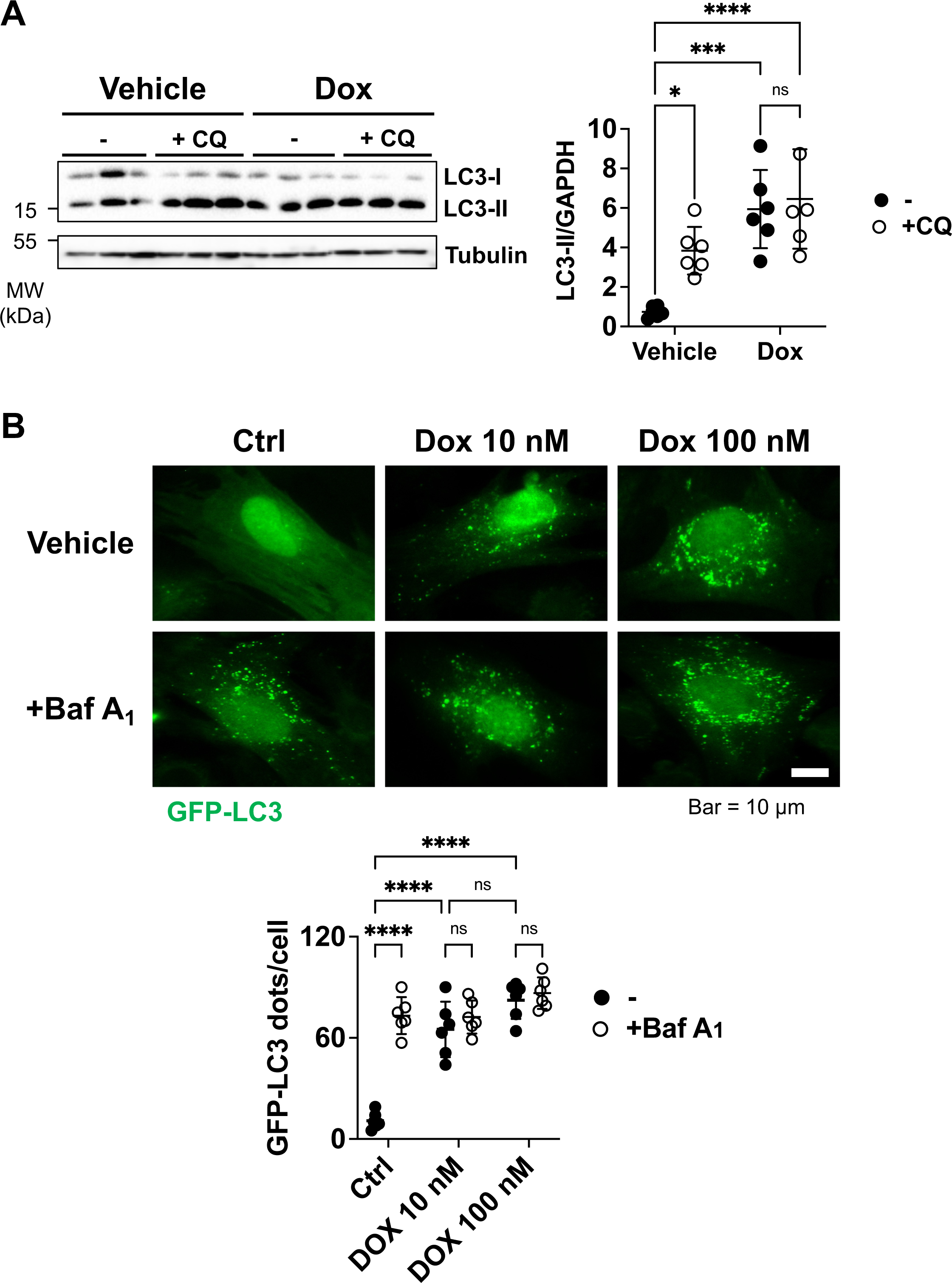
Doxorubicin (Dox) inhibited autophagic flux in cardiomyocytes. A. C57BL/6J mice were administered Dox (5 mg/kg) or vehicle by once-per-week intraperitoneal (i.p.) injection for 4 weeks. Mouse hearts were harvested 1 week after the final Dox injection. Chloroquine (CQ) (10 𝛍g/kg i.p.) was administered 4 hours before harvesting the heart. Protein levels of p62 and LC3 in the heart were analyzed by immunoblot analyses. Representative immunoblot (**left**) and summary of the ratios of p62 to GAPDH and LC3-II to GAPDH (**right**). n (the number of animals) =6 per group. One-way ANOVA with Bonferroni’s post hoc test. **** P<0.0001, *** P<0.001, * P<0.05, ns, not significant. P<0.05 was considered significant. B. Neonatal rat ventricular myocytes (NRVMs) were transduced with GFP-LC3 adenovirus with vehicle or Dox treatment for 48 hours. Bafilomycin A1 (Baf A_1_) was applied for 3 hours before cell fixation. The number of GFP-LC3 puncta per cell was counted (**right**). Representative images are shown (**left**). Scale bar = 10 𝛍m. The image shown is representative of six independent experiments. n (the number of experiments) =6 in each group. Data are presented as Mean±SEM. One-way ANOVA with Bonferroni’s post hoc test. **** P<0.0001, ns, not significant. P<0.05 was considered significant.

### 3.6 Suppression of autophagy promotes cardiomyocyte senescence and cardiac dysfunction in Dox-treated mouse hearts

In order to investigate the role of autophagy suppression in mediating senescence in the mouse model of Dox-induced cardiomyopathy, the effect of Dox was evaluated in mice with a gain of autophagy function. We have shown previously that phosphorylation of mouse Beclin 1 at Threonine 106, equivalent to human Beclin 1 Threonine 108, induces association of Beclin 1 and Bcl-2, thereby inhibiting Beclin 1 and autophagy [51]. Expression of Beclin 1(T106A), a Threonine 106 phosphorylation-resistant mutant of mouse Beclin 1, potently increases autophagic flux in cardiomyocytes [51]. We generated Beclin 1(T106A) knock-in mice and confirmed that they have normal cardiac function at baseline (Supplementary Table 4). Autophagic flux was stimulated in Beclin 1(T106A) mice (**Supplemental Fig. 5**). Dox-induced increases in γH2AX staining were abrogated in Beclin 1(T106A) knock-in mice (**Fig. 8AB**). Immunoblot analyses showed that Dox-induced upregulation of P53 and P21 is inhibited in Beclin 1(T106A) knock-in mice (**Fig. 8C**). Furthermore, Dox-induced LV chamber dilation and decreases in LVEF and %FS were completely abrogated in Beclin 1(T106A) knock-in mice (**Fig. 8DE and Supplementary Table 4**). Dox-induced decreases in cardiac fibrosis were also abrogated in Beclin 1(T106A) knock-in mice (**Fig. 8F**). These results suggest that the suppression of autophagy plays an important role in mediating cardiomyocyte senescence and cardiac dysfunction in the Dox-induced cardiomyopathy mouse model.

**Figure 8.**
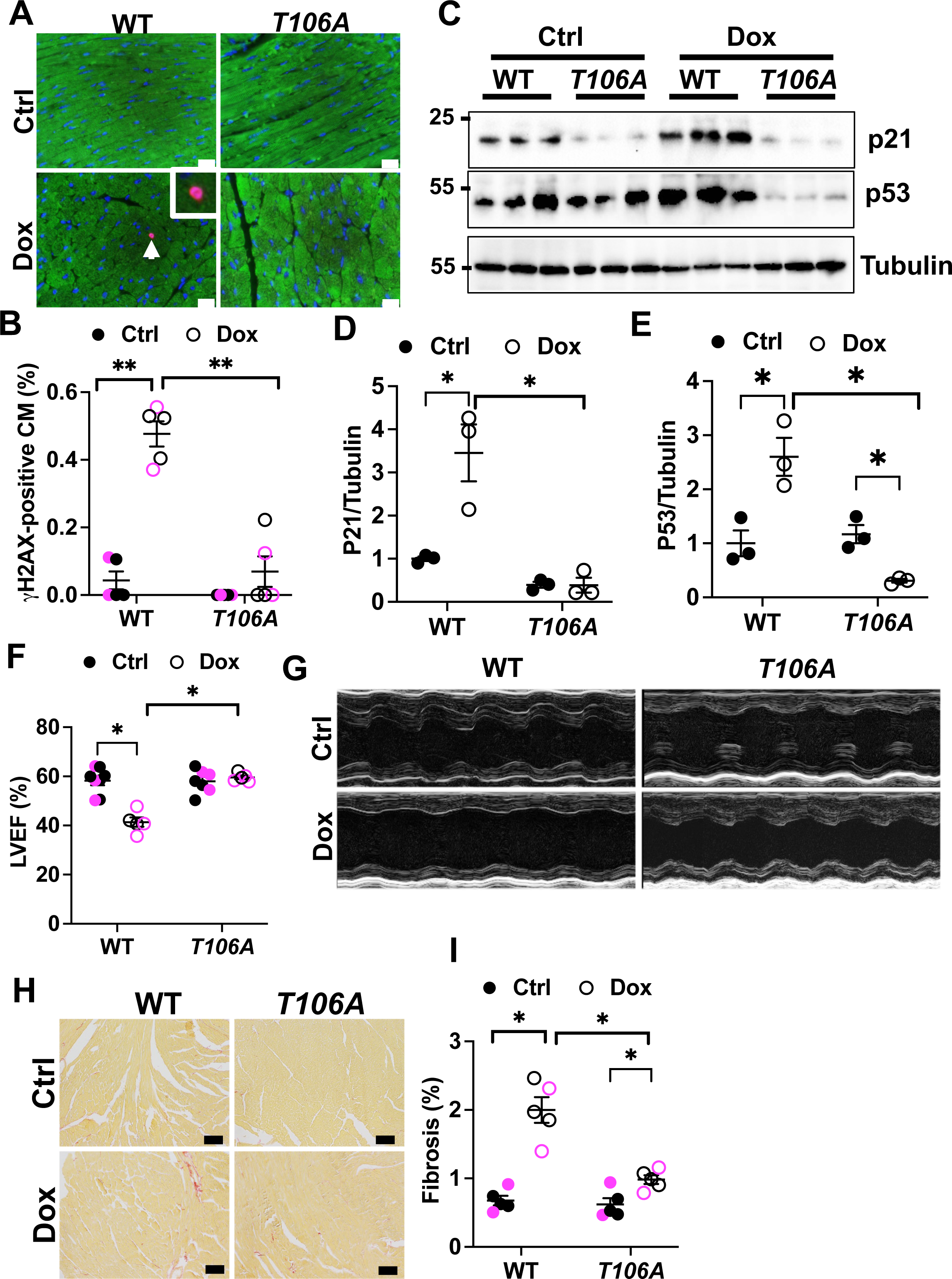
*Beclin 1*(T106A) attenuates doxorubicin-induced cardiomyopathy in mice. A. Images of γH2AX (red) and cTnt (green) immunostaining in PBS-(Ctrl) or doxorubicin (Dox)-treated wild type mice (WT) and *T106A* mice. B. Percentage of γH2AX-positive cardiomyocytes in PBS-(Ctrl) or Dox-treated WT and *T106A* mice. n (the number of animals) =6 per group. ** P<0.01. C. Immunoblots of P21, P53, and Tubulin in Ctrl and Dox-treated WT and *T106A* mice. D. Relative band density of P21. E. Relative band density of P53. In C-D, n (the number of animals) =3 per group. *P<0.05. F. Left ventricular ejection fraction (LVEF) in Dox-treated or PBS (Ctrl)-treated WT or *T106A* mice. n (the number of animals) =6 per group. * P<0.05. G. Representative M-mode echo images. H. Images of PASR staining of cardiac tissue sections from WT and *T106A* mice after Dox or Ctrl treatments. Scale bars = 100 μm. I. Percentage of fibrosis (PASR-positive area). n (the number of animals) =6 per group. * P<0.05. In B, F and I, black dots/circles show data from male mice and pink dots/circles show data from females. Data are presented as Mean±SEM. Statistical analyses were conducted using Prism 2-way ANOVA followed by multiple comparison t test and/or Bonferroni’s t test. P<0.05 was considered significant.

## 4. Discussion

Our results suggest that suppression of autophagy promotes senescence in cardiomyocytes both *in vivo* and *in vitro* through a cell-autonomous mechanism. Suppression of autophagy plays a causative role in promoting cardiomyocyte senescence in the mouse model of Dox-induced cardiomyopathy.

We found that inhibition of autophagy increases typical indicators of senescence in cardiomyocytes, including cell cycle inhibitors (P16 and P21), P53, a DNA damage marker (γH2AX) and SA-β-gal. In addition, inhibition of autophagy induces upregulation of proinflammatory cytokines, including IL-6 and TNF-α, in cardiomyocytes, consistent with the notion that inhibition of autophagy promotes the SASP. Furthermore, senolytic treatment decreased the number of γH2AX-positive cardiomyocytes in *Atg7*cKO mouse hearts. Taken together, these results suggest that suppression of autophagy stimulates senescence in cardiomyocytes.

Although the level of induction of senescence markers following Atg7 downregulation or Dox treatment is similar to that produced by other interventions to induce senescence in the heart [52], it is likely that the individual markers detect non-senescent cells as well as senescent cells. For example, P16-positive cells may consist of not only those exhibiting features of cellular senescence but also those not involved in senescence [53]. Thus, at present, it is essential to use multiple markers of senescence to define induction of senescence. Convenient methods to separate a senescent cardiomyocyte pool defined by multiple indicators need to be developed, and then further characterization, such as deep sequencing, should be conducted to better define senescent cardiomyocytes [54].

Although the role of autophagy in mediating senescence appears to be cell-type- and context-dependent [13], we show that both genetic downregulation of Atg7, which inhibits autophagosome formation, and Dox treatment, which inhibits autophagic flux, promote senescence in cardiomyocytes. Induction of senescence was observed in mice with chimeric, i.e., only partial, downregulation of Atg7. Thus, induction of cardiomyocyte senescence in the *Atg7*cKO mouse heart is independent of the presence of any cardiac dysfunction. Furthermore, induction of senescence in cardiomyocytes was also observed when Atg7 was downregulated in NRVMs or when NRVMs were treated with Dox. Collectively, these results suggest that downregulation of autophagy can directly stimulate cardiomyocyte senescence via cell-autonomous mechanisms.

What is the underlying cellular mechanism through which downregulation of autophagy induces senescence in cardiomyocytes? Since cardiomyocytes in adult mouse hearts rarely divide, it is unlikely that induction of senescence in *Atg7*cKO mouse hearts and Dox-induced cardiomyopathy is mediated through cardiomyocyte proliferation and telomere shortening. It has been shown that cardiomyocytes can undergo senescence in a telomere length-independent manner and through stimulation of telomere damage caused by oxidative stress [19]. There are several possible mechanisms. First, downregulation of autophagy is accompanied by increases in γH2AX-positive cardiomyocytes. Downregulation of autophagy could induce accumulation of misfolded and dysfunctional proteins and organelles [55]. The resulting increases in cellular stress may, in turn, induce oxidative stress and DNA damage. Second, increased expression of proinflammatory cytokines increases oxidative stress and cell death, which may, in turn, facilitate inflammasome-mediated cell signaling, thereby facilitating senescence. Finally, autophagy may directly affect cell signaling through modulation of the stability of signaling molecules. For example, we have shown recently that YAP, a transcription factor co-factor and the terminal effector of the Hippo pathway, is degraded by autophagy under baseline conditions and, thus, downregulation of autophagy induces accumulation of YAP in cardiomyocytes [56]. Further investigation is warranted to clarify the molecular mechanism by which downregulation of autophagy stimulates senescence in cardiomyocytes.

The SASP is a major mechanism through which senescent cells propagate their actions to surrounding cells through autocrine/paracrine mechanisms and affect the pathophysiology of organs and even whole organisms [57]. Screening with cytokine array analyses showed that Atg7 downregulation in cardiomyocytes not only upregulates proinflammatory cytokines, including CINC-2a, IL-6, B7-2/CD86, RAGE and TNF-α, but also downregulates several cytokines known to possess cell protective attributes, including CNTF, IL-1R6, IL-2 and CXCL5 [19]. Further investigation is needed to clarify the functional significance of the changes in the level of the aforementioned factors in the heart. It would also be interesting to clarify the molecular mechanism through which autophagy suppression affects the levels of these cytokines in the heart.

ABT-263 is a potent inhibitor of the Bcl-2 family proteins and induces apoptosis of senescent cells [38, 49]. We observed that the number of γH2AX-positive cardiomyocytes in *Atg7*cKO mice is decreased in response to ABT-263 treatment. We speculate that induction of apoptosis in senescent cells is followed by rapid elimination by efferocytosis. Further investigation is needed to clarify the mechanism mediating the death of senescent cardiomyocytes in the presence of ABT-263. It should be noted that we cannot exclude the possibility that ABT-263 treatment also induces senolysis in non-cardiomyocytes. ABT-263 treatment significantly improved cardiac function in *Atg7*cKO mice, Thus, our result suggests that increased senescence in cardiomyocytes may play an important role in mediating the development of cardiomyopathy in *Atg7*cKO mice.

Aside from senolysis, which eliminates cells that have already become senescent, an intervention that prevents cells from becoming senescent is also effective in protecting the heart against Dox-induced cardiomyopathy. We show here that Beclin 1(T106A) knock-in mice, which exhibit increased autophagy, exhibited reduced senescence and improved cardiac function even in the presence of Dox treatment. These results suggest that autophagy suppression plays an important role in mediating increases in cardiomyocyte senescence and the development of cardiomyopathy in response to Dox treatment.

Interestingly, the intervention to *eliminate senescent cardiomyocytes* and the intervention to *promote survival of non-senescent cardiomyocytes* are both effective in preventing the development of cardiomyopathy. Thus, two different cardiomyocyte populations in the heart, namely senescent cardiomyocytes and non-senescent cardiomyocytes, could be targeted to regulate their survival in opposite directions, namely inhibiting survival of senescent cardiomyocytes and promoting survival of non-senescent cardiomyocytes. Although the conventional therapy for heart failure focuses on preventing the death of functional cardiomyocytes, it may be necessary to add additional interventions to promote the death of senescent cardiomyocytes. If true, it may also be necessary to determine why it would be beneficial to eliminate senescent cardiomyocytes even though they retain the ability to contract.

We acknowledge limitations in this study. First, whether or not the downregulation of autophagy by aging has a causative role in mediating senescence in cardiomyocytes requires further testing, using aged mice. It will be interesting to determine whether aging in *Atg7*cKO mice has an additive effect upon cardiomyocyte senescence and the development of cardiomyopathy. Since both *Atg7*cKO and aging inhibit autophagy, Atg7 downregulation and aging may not exhibit additive effects. Second, some experiments were compromised by the small number of mice, which was caused by inefficient breeding of the mouse lines for unknown reasons.

In summary, we here demonstrated that downregulation of autophagy induces senescence in cardiomyocytes. Autophagy is inhibited in many conditions in the heart, including aging, obesity cardiomyopathy, and during the chronic phase of heart failure [55]. Whether suppression of autophagy plays a causative role in senescence in these conditions and, if so, whether senescence is involved in the development of cardiomyopathy remains to be investigated.

## Author contributions

P.Z. and J.S. designed the experiments and wrote the paper; P.Z., E.-A. S. and Y.T. conducted the *in vitro* and *in vivo* experiments; K.T. conducted echocardiographic measurements of mice. Y.T., E.-A. S., Y.S.-W.. and P.Z. conducted the animal experiments and analyses; E.-A. S. and Y.T. conducted immunohistochemistry analyses; J.S. generated project resources. All authors reviewed and commented on the manuscript.

## Data and material availability

The data that support findings, including statistical analyses, and reagents used are available from the corresponding author upon request.

## Acknowledgement

We thank Daniela Zablocki for critical reading of the manuscript.

## Supplementary data

Supplementary data to this article can be found online at https://XX

## Abbreviations

SASP: (senescence-associated secretory phenotype)
Atg7cKO: (cardiac-specific Atg7 knockout)
CM: (Cardiomyocyte)
WT: (wild type)
Dox: (doxorubicin)
AAV9: (adeno-associated virus 9)
cTNT: (cardiac troponin T)
GFP: (green fluorescence protein)
LV: (left ventricle/ventricular)
LVESD: (left ventricular end-systolic dimension)
LVEF: (left ventricular ejection fraction)
LVFS: (left ventricular fractional shortening)
LVEDD: (left ventricular end-diastolic dimension)
CQ: (chloroquine)
Baf A1: (Bafilomycin A1)
SA-βgal: (senescence-associated β-galactosidase)
LC3: (Microtubule-associated protein 1A/1B-light chain 3)
DAPI: (4’,6-diamidino-2-phenylindole)
IL-6: (interleukin 6)
NRVM: (neonatal rat ventricular myocyte)
Ctrl: (Control)
H2AX: (H2A histone family member X)
Yap: (yes-associated protein)
KO: (knockout)
IL-1: (interleukin 1)
IL-10: (interleukin 10)
CINC-2a: (cytokine-induced neutrophil chemoattractant-2alpha)
MIP-3a: (Macrophage Inflammatory Protein-3 alpha)
RAGE: (Receptor for advanced glycation end products)
TNF-α: (Tumor necrosis factor alpha)
CNTF: (Ciliary neurotrophic factor)
IL-1R6: (interleukin-36 receptor)
CXCL5: (C-X-C motif chemokine ligand 5)
IL-2: (interleukin 2)
LIX: (Lipopolysaccharides-induced CXC chemokine)

---

**Supplementary Table 1.** Echocardiographic measurements of *Atg7flox/flox* mice infected with AAV9-*cTnt-Cre-Gfp*.

|  | Ctrl | *cTnt-Cre-Gfp* |
| --- | --- | --- |
| n | 3 | 3 |
| IVSd (mm) | 0.70±0.02 | 0.62±0.04 |
| LVIDd (mm) | 4.10±0.25 | 4.47±0.06 |
| LVIDs (mm) | 2.87±0.22 | 3.32±0.06 |
| LVPWd (mm) | 0.67±0.01 | 0.62±0.03 |
| EF (%) | 61±1 | 54±1 <sup>A</sup> |
| FS (%) | 27±1 | 23±1 <sup>A</sup> |
| HR (bpm) | 476±8 | 481±5 |

A p<0.01 vs. Ctrl.
IVSd interventricular septal end diastole dimension; LVEDD left ventricular end-diastolic diameter; LVESD left ventricular-systolic diameter; LVPWd left ventricular posterior wall end diastole dimension; EF ejection fraction; FS fractional shortening; HR heart rate.

**Supplementary Table 2.** Cytokines regulated by Atg7 knockdown in cardiomyocytes.

| Cytokines upregulated by shAtg7 | Fold | Cytokines downregulated by shAtg7 | Fold |
| --- | --- | --- | --- |
| CINC-2a | 1.63 | CNTF | 0.72 |
| B7-2/CD86 | 1.58 | IL-1R6 | 0.56 |
| MIP-3a | 1.64 | IL-2 | 0.72 |
| RAGE | 1.51 | LIX (CXCL5) | 0.77 |
| TNF-α | 1.52 | PDGF-AA | 0.82 |
| CINC-1 | 1.21 | Thymus chemokine-1 | 0.78 |
| Agrin | 1.21 | IL-13 | 0.73 |
| Activin-A | 1.17 | GMCSF | 0.97 |
| β-NGF | 1.15 | TIMP1 | 0.90 |
| CINC-3 | 1.23 |  |  |
| IL-1a | 1.13 |  |  |
| IL-1b | 1.15 |  |  |
| Prolactin R | 1.32 |  |  |
| Fas Ligand | 1.21 |  |  |
| ICAM-1 | 1.08 |  |  |
| IFN-gamma | 1.04 |  |  |
| IL-4 | 1.19 |  |  |
| IL-10 | 1.09 |  |  |
| L-selectin | 1.01 |  |  |
| MCP-1 | 1.05 |  |  |
| MMP-8 | 1.09 |  |  |
| VEGF | 1.05 |  |  |
| Leptin | 1.02 |  |  |

**Supplementary Table 3.**
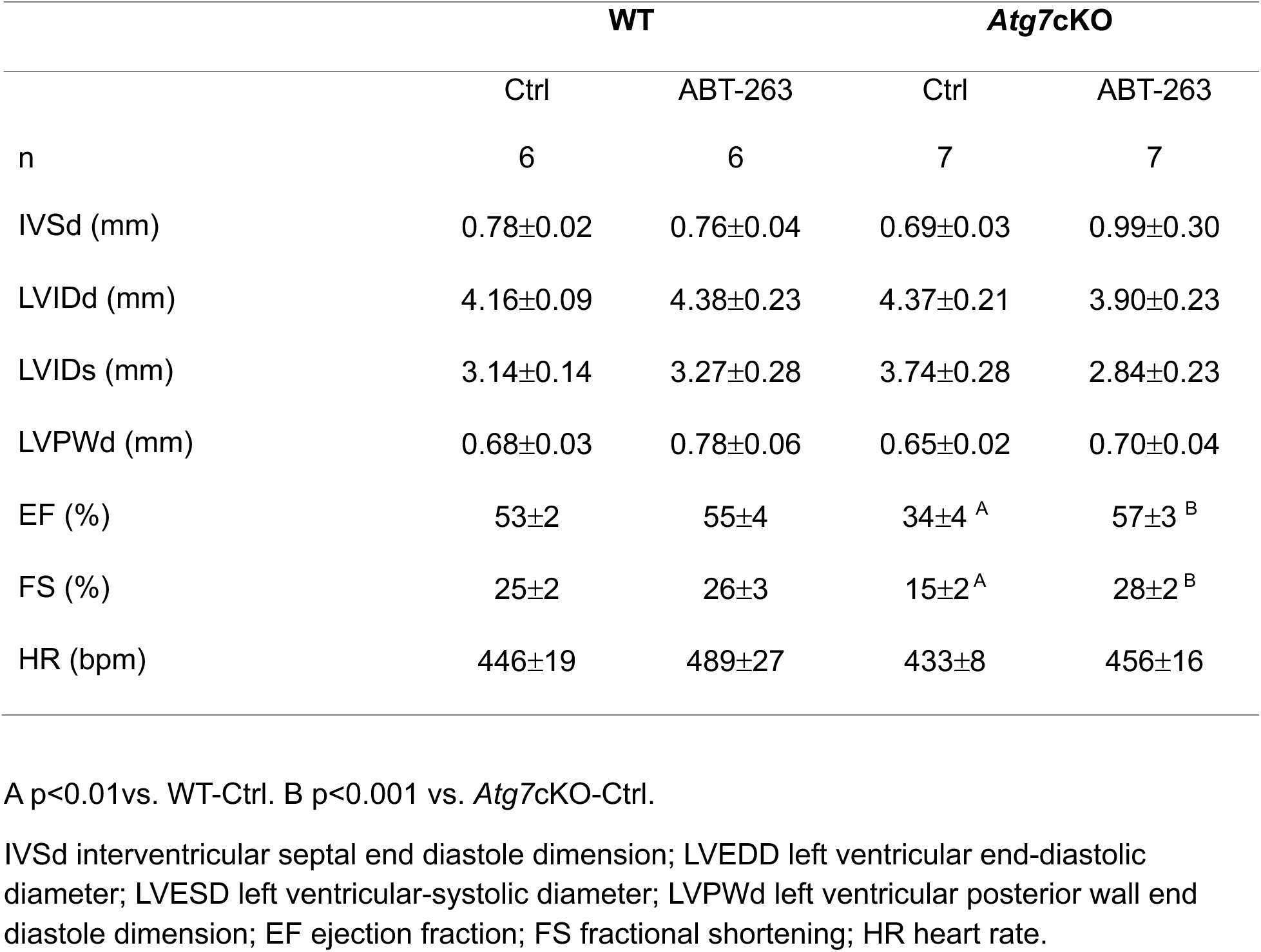
Echocardiographic measurements of *Atg7*cKO mice treated with ABT-263.

|  | WT |  | *Atg7cKO* |  |
| --- | --- | --- | --- | --- |
|  | Ctrl | ABT-263 | Ctrl | ABT-263 |
| n | 6 | 6 | 7 | 7 |
| IVSd (mm) | 0.78±0.02 | 0.76±0.04 | 0.69±0.03 | 0.99±0.30 |
| LVIDd (mm) | 4.16±0.09 | 4.38±0.23 | 4.37±0.21 | 3.90±0.23 |
| LVIDs (mm) | 3.14±0.14 | 3.27±0.28 | 3.74±0.28 | 2.84±0.23 |
| LVPWd (mm) | 0.68±0.03 | 0.78±0.06 | 0.65±0.02 | 0.70±0.04 |
| EF (%) | 53±2 | 55±4 | 34±4 <sup>A</sup> | 57±3 <sup>B</sup> |
| FS (%) | 25±2 | 26±3 | 15±2 <sup>A</sup> | 28±2 <sup>B</sup> |
| HR (bpm) | 446±19 | 489±27 | 433±8 | 456±16 |

A p<0.01vs. WT-Ctrl. B p<0.001 vs. *Atg7cKO*-Ctrl.
IVSd interventricular septal end diastole dimension; LVEDD left ventricular end-diastolic diameter; LVESD left ventricular-systolic diameter; LVPWd left ventricular posterior wall end diastole dimension; EF ejection fraction; FS fractional shortening; HR heart rate.

**Supplementary Table 4.** Echocardiographic measurements of *Beclin1(T106A)* knockin mice treated with Dox.

|  | **WT** |  | ***Beclin1(T106A)*** |  |
| --- | --- | --- | --- | --- |
|  | Ctrl | Dox | Ctrl | Dox |
| n | 8 | 5 | 7 | 5 |
| IVSd (mm) | 0.72±0.04 | 0.77±0.13 | 0.66±0.05 | 0.77±0.08 |
| LVIDd (mm) | 3.68±0.10 | 4.63±0.21<sup>A</sup> | 3.98±0.14 | 4.29±0.23 |
| LVIDs (mm) | 2.62±0.11 | 3.80±0.17<sup>A</sup> | 2.80±0.15 | 3.08±0.21<sup>C</sup> |
| LVPWd (mm) | 0.62±0.02 | 0.69±0.11 | 0.57±0.02 | 0.62±0.06 |
| EF (%) | 58±2 | 41±2<sup>B</sup> | 58±2 | 60±2<sup>D</sup> |
| FS (%) | 29±1 | 18±1<sup>A</sup> | 30±2 | 28±1<sup>D</sup> |
| HR (bpm) | 429±17 | 464±20 | 432±11 | 464±25 |

A p<0.01, B p<0.001 vs. WT-Ctrl. C p<0.05, D p<0.001 vs. WT-Dox.
IVSd interventricular septal end diastole dimension; LVEDD left ventricular end-diastolic diameter; LVESD left ventricular-systolic diameter; LVPWd left ventricular posterior wall end diastole dimension; EF ejection fraction; FS fractional shortening; HR heart rate.

**Supplementary Figure 1.**
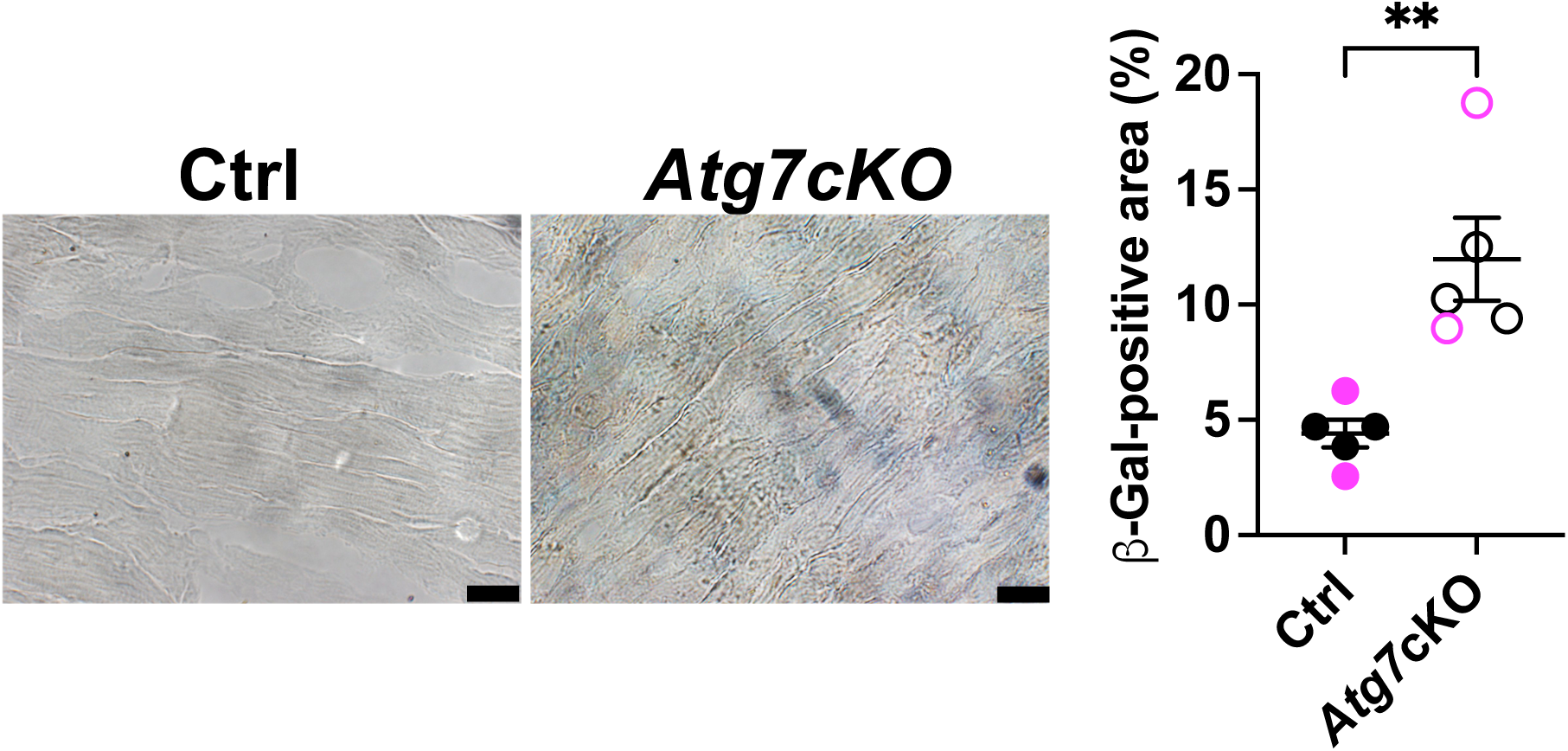
The *Atg7* cKO mouse heart exhibit more senescence. The heart sections were subjected to β-galactosidase staining. Scale bars = 20 μm. **p<0.01. n=5 in each group.

**Supplementary Figure 2.**
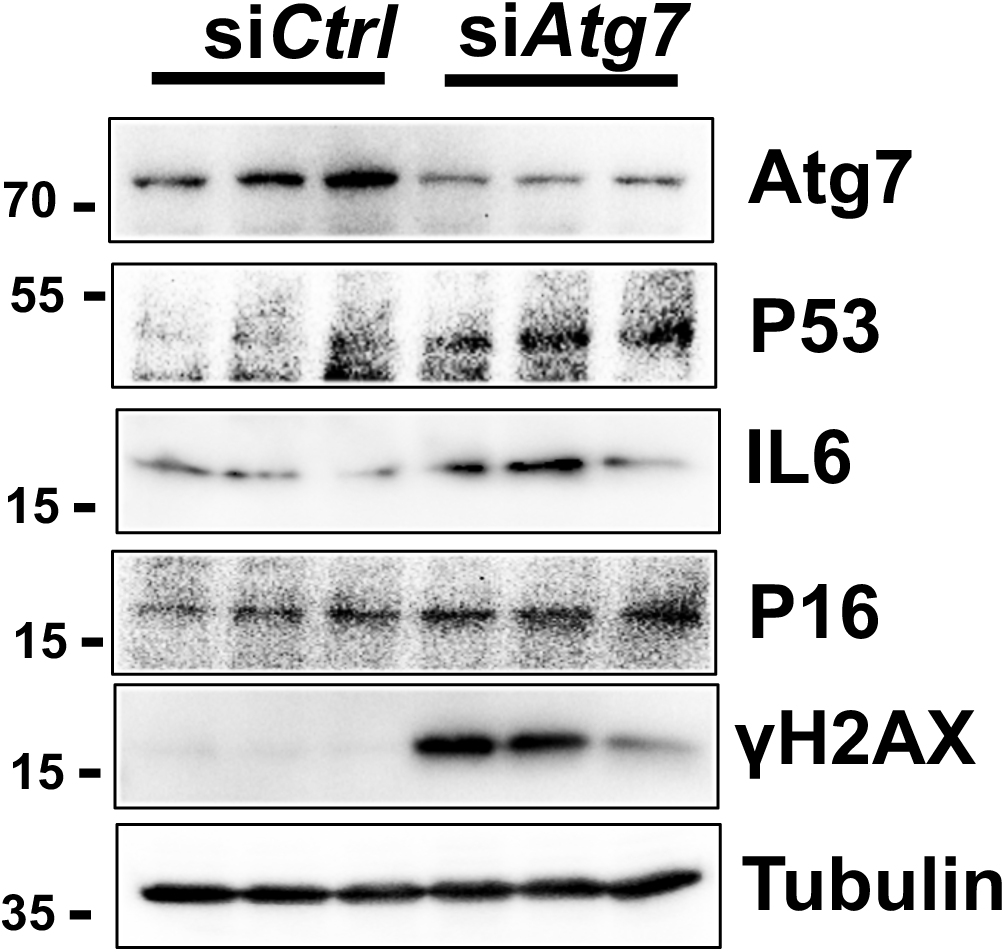
Knockdown of *Atg7* promotes senescence in cultured cardiomyocytes. Neonatal rat ventricular cardiomyocytes were treated with siRNA control (siCtrl) or siRNA Atg7 (siAtg7). Immunoblot analyses of Atg7, P53, IL6, p16, γH2AX, and Tubulin are shown.

**Supplementary Figure 3.**
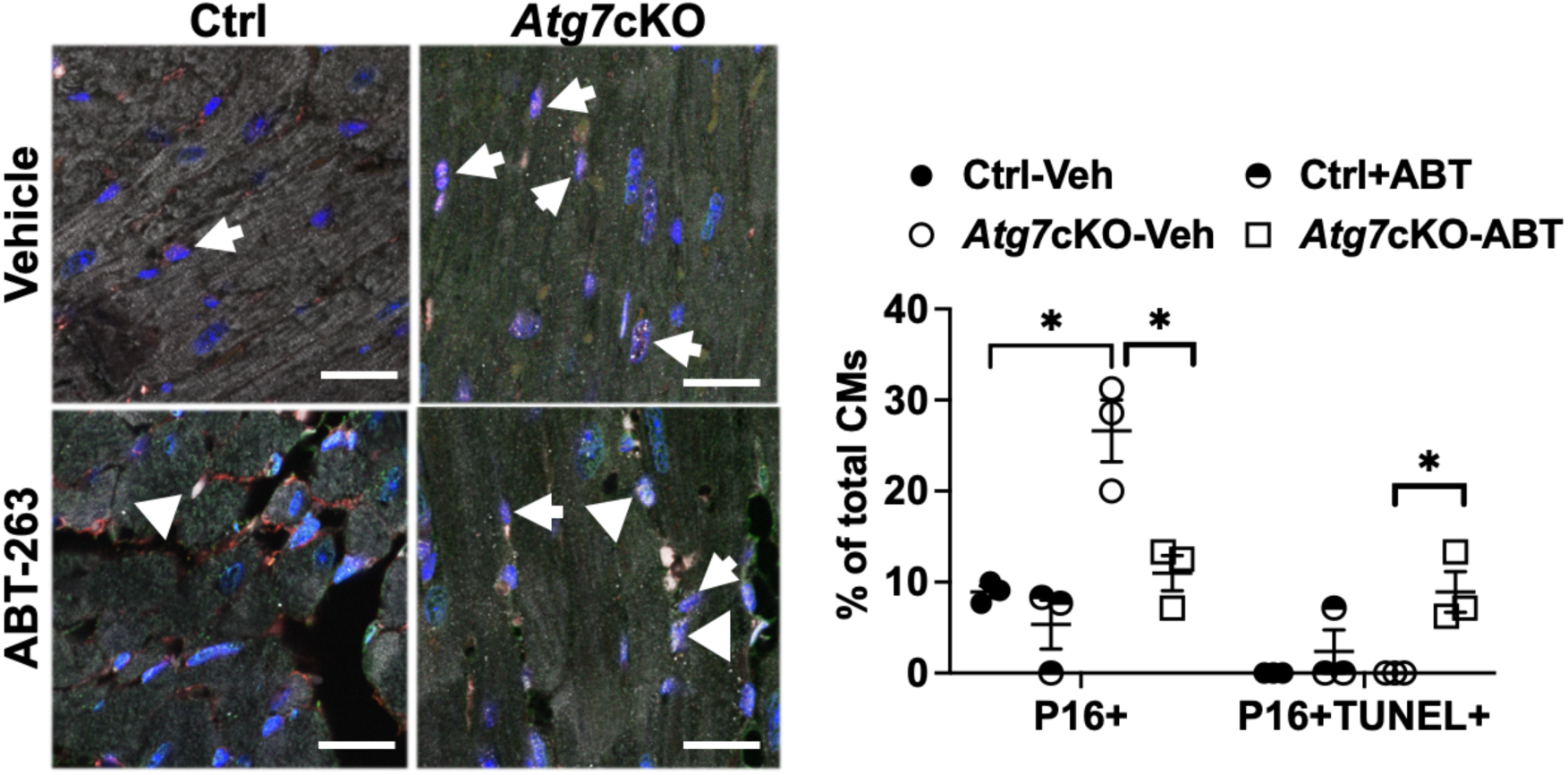
Control (Ctrl) and *Atg7*cKO mice were treated with either vehicle or ABT-263 as indicated in Fig. 5A. Myocardial sections were subjected to TUNEL staining and then stained with anti-P16 antibody and anti-cTNT antibody, and lastly stained with DAPI. The number of P16-and cTNT-double positive cells and that of P16-(red), cTNT-(white) and TUNEL-(green) triple positive cells was quantitated. *p<0.05. White arrows indicate P16-positive nuclei in cTNT-positive cells (cardiomyocytes) whereas white triangles indicate P16- and TUNEL-double positive nuclei in cTNT-positive cells (cardiomyocytes). Scale bars = 20 µm. N=3 in each group.

**Supplementary Figure 4.**
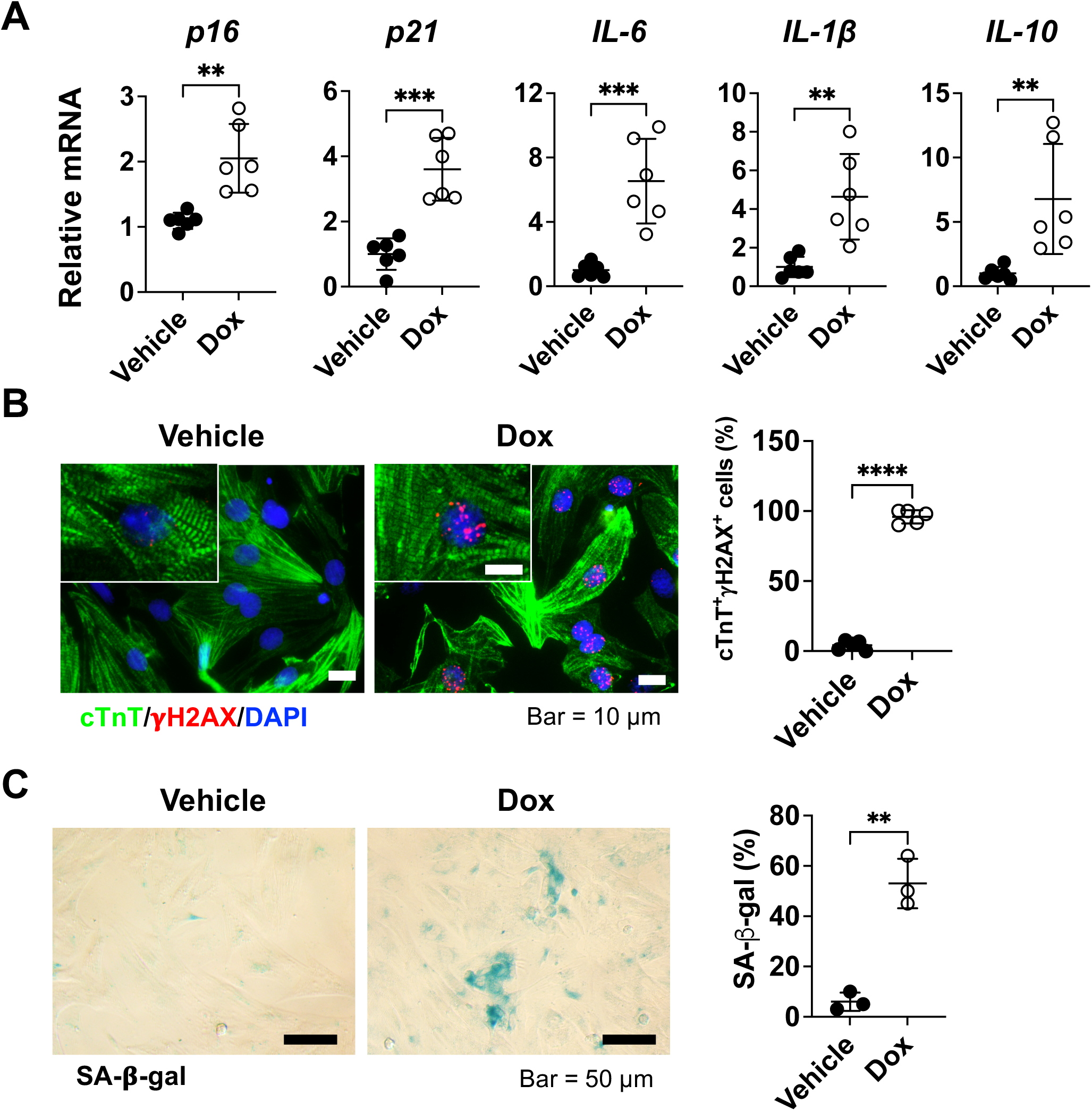
A. *p16*, *p21*, *IL-6*, *IL-1β*, and *IL-10* mRNA expression levels in vehicle- or Dox-treated mouse hearts. In B-C, n (the number of animals) =6. ** P< 0.01, *** P< 0.001. B. Neonatal rat ventricular myocytes (NRVMs) were treated with vehicle or Dox (100 nM) for 48 hours and analyzed by immunofluorescence staining. The number of γH2AX^+^cTnT^+^ cells was counted and is shown in the dot plots. Scale bar = 10 μm. **** P<0.0001. n (the number of experiments) =6. C. After Dox treatment for 48 hours, NRVMs were analyzed by SA-β-gal staining. Scale bar = 50 μm. The dot plot shows the percentage of positive SA-β-gal staining. n (the number of experiments) =3.

**Supplementary Figure 5.**
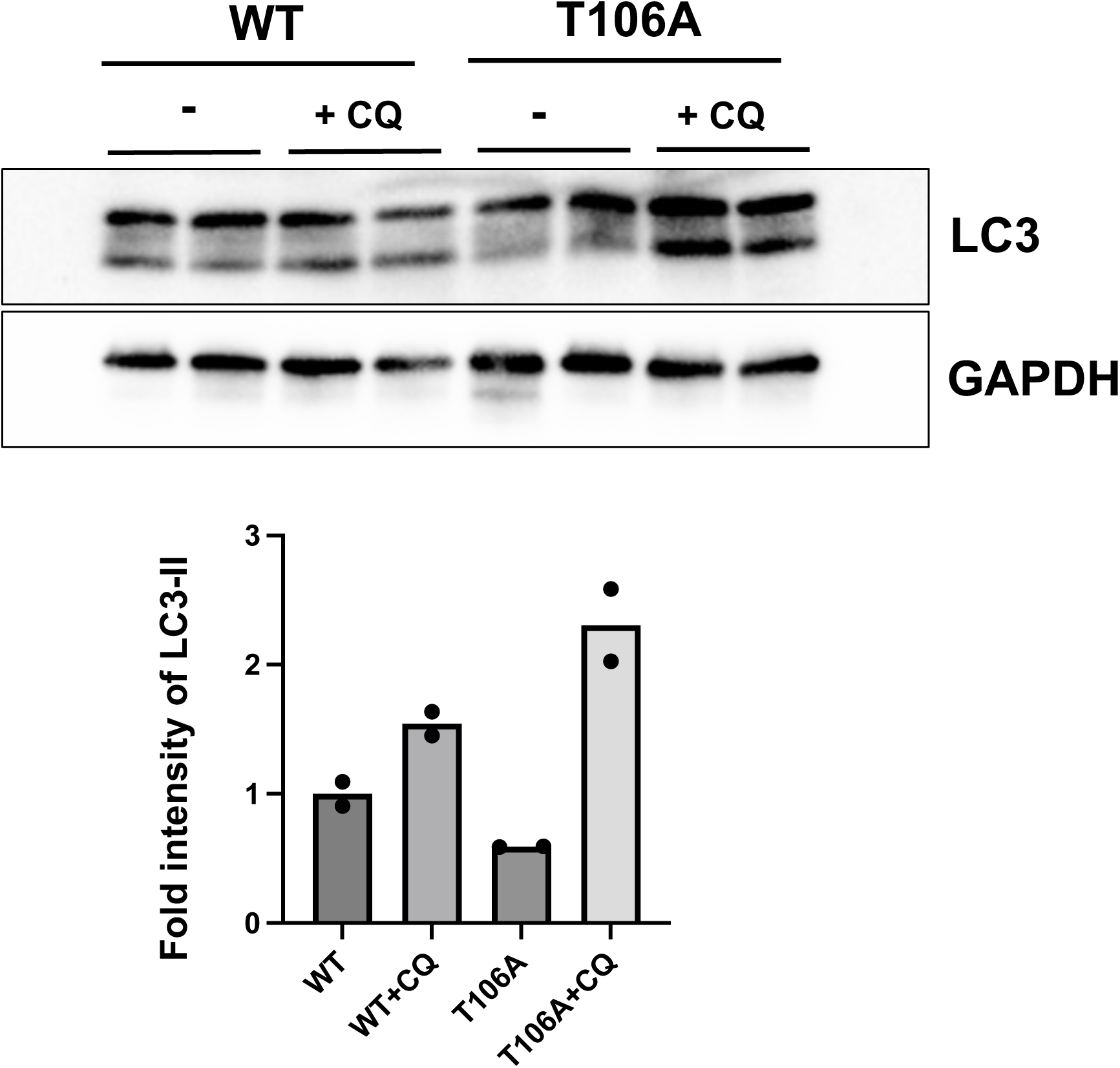
Autophagic flux was stimulated in Beclin1 (T106A) mice. Wild type and Beclin1 (T106A) mice were subjected to either saline or chloroquine injection (10 μg/kg). After 3 hours, the heart was harvested and immunoblot analyses were conducted. The graph shows the intensity of LC3II normalized by that of GAPDH. N=2 per group.

